# A mutational hotspot that determines highly repeatable evolution can be built and broken by silent genetic changes

**DOI:** 10.1101/2021.01.04.425178

**Authors:** James S. Horton, Louise M. Flanagan, Robert W. Jackson, Nicholas K. Priest, Tiffany B. Taylor

**Affiliations:** Milner Centre for Evolution, Department of Biology & Biochemistry, University of Bath, Claverton Down, Bath BA2 7AY, UK; School of Biosciences and Birmingham Institute of Forest Research (BIFoR), University of Birmingham, Edgbaston, Birmingham, B15 2TT, UK

**Keywords:** Repeatable evolution, Mutational hotspot, Silent mutation, Mutation rate heterogeneity, Experimental evolution

## Abstract

Mutational hotspots can determine evolutionary outcomes and make evolution repeatable. Hotspots are products of multiple evolutionary forces including mutation rate heterogeneity, but this variable is often hard to identify. In this work we reveal that a powerfully deterministic genetic hotspot can be built and broken by a handful of silent mutations. We observed this when studying homologous immotile variants of the bacteria *Pseudomonas fluorescens*, AR2 and Pf0-2x. AR2 resurrects motility through highly repeatable *de novo* mutation of the same nucleotide in >95% lines in minimal media (*ntrB* A289C). Pf0-2x, however, evolves via a number of mutations meaning the two strains diverge significantly during adaptation. We determined that this evolutionary disparity was owed to just 6 synonymous variations within the *ntrB* locus, which we demonstrated by swapping the sites and observing that we were able to both break (>95% to 0% in AR2) and build (0% to 80% in Pf0-2x) a powerfully deterministic mutational hotspot. Our work reveals a fundamental role for silent genetic variation in determining adaptive outcomes.

## Introduction

Mutational hotspots, which describe instances where independent cell lines persistently fix mutations at the same genomic sites, can make evolution remarkably repeatable. Such hotspots are of immense importance as they have been observed to drive evolution across the domains of life, from viruses (including SARS-CoV-2; Weber et al. 2020), to bacteria (including MRSA; Sekowska et al. 2016), to higher eukaryotic cell lines including those in avian species (Galen et al. 2015) and human cancers (Trevino 2020). Our understanding of evolutionary dynamics (e.g. competitive selection and clonal interference) can sometimes explain the appearance of hotspots, but genetic features that build hotspots by biasing mutation rates are much less understood.

There have been many examples of experimental systems evolving via repeatable evolution. Microbes evolving under strong selection often rapidly adopt similar novel phenotypes (Fong et al. 2005; Ostrowski et al. 2008). Furthermore, these phenotypes are often underpinned by mutation hotspots, which come in the form of clustered genetic changes within the same region of the genome (Riehle et al. 2001; Fraebel et al. 2017), or within limited pockets of loci (Bull et al. 1997; Wichman et al. 1999; Herron and Doebeli 2013; Kram et al. 2017). Sometimes realised mutations are found only in genes from a single regulatory pathway (Notley-McRobb and Ferenci 1999; Miller et al. 2013) or a single protein complex (Avrani et al. 2017). In extreme cases, evolutionary events can be seen to repeatedly target just a handful of sites within a single locus (Meyer *et al*., 2012; van Ditmarsch *et al*., 2013). Repeatable evolution allows lines to evolve in parallel, and the degree of parallelism typically becomes less common as it descends from broader genomic regions to the nucleotide (Tenaillon et al. 2012; Bailey et al. 2015). However, despite frequent descriptions of repeatable evolutionary events, a detailed understanding of the hotspots that ensure their occurrence is often lacking.

There are three primary facilitators of mutational hotspots that drive repeatable evolution: (*i*) Fixation bias, which skews evolution toward mutations that enjoy a higher likelihood of dominating the population pool. Not all facilitators of fixation bias are considered adaptively advantageous (Eyre-Walker and Hurst 2001), but in instances where we observe rapid and highly parallel sweeps it will likely take the form of selection, which drives the fittest competing genotypes in the population to fixation (see Wood, Burke and Rieseberg, 2005; Woods *et al*., 2006). (*ii*) Mutational accessibility, as there may be only a small number of readily accessible mutations a genotype can undergo to improve fitness (Weinreich et al. 2006). And, (*iii*) Mutation rate heterogeneity, where genetic and molecular features scattered throughout the genome cause sites to radiate at different rates, introducing a mutation bias toward a particular outcome (Bailey et al. 2017). Previous research shows that mutation rate heterogeneity can be influenced by the arrangement of nucleotides surrounding a particular site (Long et al. 2014), and genetic features such as the secondary structure of DNA (Duan et al. 2018) including the formation of single-stranded DNA hairpins (De Boer and Ripley 1984). Nevertheless, the prominence of genetic sequence in driving parallel evolutionary outcomes remains unknown.

To establish which mechanisms are at play, it is important to consider whether parallel outcomes are robust to experimental conditions such as environment (Turner et al. 2018) and to account for clonal interference, which can alter the chance of observing parallel evolution (Bailey et al. 2017; Lässig et al. 2017). Clonal interference can occur either due to standing genetic variation in the founder population which yields multiple adaptive genotypes in a novel environment (i.e. a soft selective sweep; Hermisson and Pennings, 2005) or when mutation rate is high relative to the selective coefficient (Barrett et al. 2006). However, clonal interference does not often play an important role when founding experimental lines with clonal samples, performing experimental procedures over short timescales, and ensuring rapid fixation of adaptive mutants e.g. through spatial separation and/or introducing an artificial bottleneck.

In this work, we have utilised an ideal system for identifying the key features that build mutational hotspots. We have employed two engineered non-flagellate and biosurfactant-deficient strains of the soil bacteria *P. fluorescens*: AR2, derived from SBW25, and Pf0-2x, derived from Pf0-1 (see materials and methods). The strains share homologous genetic backgrounds, including highly similar gene regulatory architectures and translated protein products, yet they evolve divergently due to local genetic differences. Both engineered strains lack function of the master regulator of flagella-dependent motility, FleQ, and both AR2 and Pf0-2x rapidly re-evolved flagella-mediated motility under strong directional selection (Taylor et al. 2015). In AR2, this phenotype was achieved in independent lineages via repeatable *de novo* mutation in the *ntrB* locus of the nitrogen regulatory (ntr) pathway. The parallel evolution of *ntrB* mutants was noteworthy as the locus was consistently targeted, whereas Pf0-2x lines evolved motility via mutations across the ntr regulatory hierarchy (Taylor et al. 2015). As such parallel evolution between these homologs varied across scale; both were parallel to the phenotype and targeted gene regulatory network, but only one possessed a mutational hotspot that concentrated mutations at a single nucleotide site within a single locus. We conducted a series of experiments to find out why.

Here we show that motility evolves in AR2 in an extremely repeatable manner, which is absent in Pf0-2x due to a genetic feature predicated on synonymous variation. The evolution of flagella motility in AR2 was found to target the same nucleotide substitution in over 95% of cases in minimal medium (M9). This outcome was found to be robust across multiple nutrient regimes both in the immotile SBW25 variant (AR2) and another SBW25 variant that was able to access biosurfactant-mediated motility prior to evolution (SBW25 Δ*fleQ*). The role of selection and the number of viable mutational routes in ensuring the parallel outcome were found to provide some explanation for parallel evolution to the level of the *ntrB* locus, but not the nucleotide. This therefore implied that intra-locus mutation rate heterogeneity was playing a critical role. We then genetically augmented the *ntrB* locus to indirectly incriminate mutation bias and revealed a key underlying genetic driver of parallel evolution. Six silent nucleotide changes were introduced within the local region around the frequently targeted site to make AR2’s genetic sequence match Pf0-2x, but without altering the protein product. These changes were found to reduce parallel evolution at the mutational hotspot from >95% to 0%. In a reciprocal experiment, silent changes introduced to the homologous strain Pf0-2x to match AR2’s local native sequence raised parallel evolution at this site from 0% to 80%. These results reveal that synonymous genetic sequence can play a dominant role in ensuring parallel evolutionary outcomes, and shines a spotlight on the overlooked mechanistic drivers behind mutational hotspots.

## Results

### SBW25-derived immotile strains evolve motility via highly repeatable evolution

To quantify the degree of parallel evolution of flagellar motility within the immotile SBW25 model system, we placed 24 independent replicates of AR2 under strong directional selection in a minimal medium environment (M9). Motile mutants were readily identified through emergent motile zones that migrated outward in a concentric circle (fig. 1A). Clonal samples were isolated from the zone’s leading edge within 24 h of emergence and their genotypes analysed through either whole-genome or targeted Sanger sequencing of the *ntrB* locus. Motile strains evolved rapidly (fig. 1B) and each independent line was found to be a product of a one-step *de novo* mutation. All 24 lines had evolved in parallel at the locus level: each had acquired a single, motility-restoring mutation within *ntrB* (fig. 1C). More surprising however, was the level of parallel evolution within the locus. 23/24 replicates had acquired a single nucleotide polymorphism at site 289, resulting in a transversion mutation from A to C (hereafter referred to as *ntrB* A289C). This resulted in a T97P missense mutation within NtrB’s PAS domain. The remaining sample had acquired a 12-base-pair deletion from nucleotide sites 406-417 (Δ406-417), resulting in an in-frame deletion of residues 136-139 (ΔLVRG) within NtrB’s phospho-acceptor domain.

**Fig. 1.**
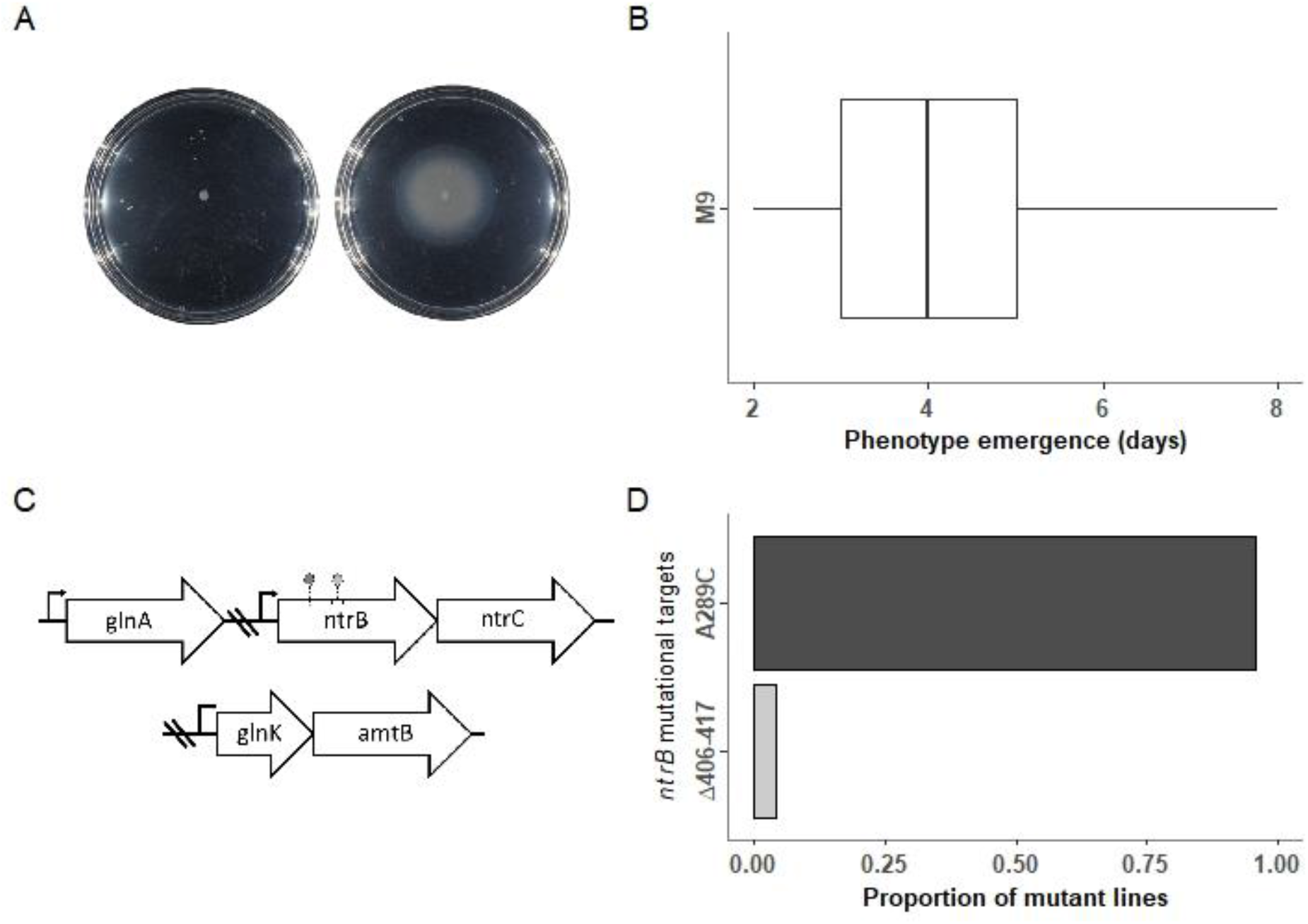
Highly repeatable evolution of flagella-mediated motility in immotile variants of *P. fluorescens* SBW25 (AR2). (A) Immotile populations evolved on soft agar (left) re-evolved flagella-mediated motility through one-step *de novo* mutation (right). (B) Phenotype emergence appeared rapidly, typically within 3-5 days following inoculation (box edges represent the 25^th^ and 75^th^ percentiles and the whiskers show the observed range). (C) The underlying genetic changes were highly parallel, with all independent lines targeting one of two sites (left circle, A289C and right circle Δ406-417) within the *ntrB* locus at the expense of other sites within the nitrogen (ntr) pathway. (D) A single transversion mutation, A289C, was the most common mutational route, appearing in over 95% of independent lines (23/24).

### Repeatable evolution is robust to nutritional environment

Repeatable evolution could be robust or highly context-dependent, especially when it occurs via *de novo* mutations with antagonistic pleiotropic effects (McGrath et al. 2011; Mcgee et al. 2016; Sackman et al. 2017). However, we found that the repeatability of the *ntrB* A289C mutation was robust across all tested conditions, despite evidence of antagonistic pleiotropic effects on growth. We tested for environment-specific antagonistic pleiotropy by measuring relative growth of the ancestral line and both evolved *ntrB* mutants on rich lysogeny broth and minimal medium containing either ammonia as the sole nitrogen source or supplemented with either glutamate (M9+glu) or glutamine (M9+gln), both of which are naturally assimilated and metabolised by the ntr system. Though large fitness costs were evident in M9 minimal medium, supplementing M9 with glu or gln reduced levels of antagonistic pleiotropy for both the *ntrB* A289C and the Δ406-417 mutants (supplementary fig. S1). Indeed, the antagonistic pleiotropy of impaired metabolism was sufficiently low in M9 supplemented with the amino acid glutamine (M9+gln) that motile mutants had increased fitness over the ancestral line in static broth, which was significant in *ntrB* A289C (*P* = 0.0361, supplementary fig. S1). These findings show that antagonistic pleiotropy is harsh in M9 and alleviated substantially in other nutritional environments, and therefore evolution in minimal media may have been limiting the viable number of adaptive routes.

We then tested whether repeatable evolution was robust to varying levels of antagonistic pleiotropy in our model system. Our expectation was that supplemented nutrient regimes would lower pleiotropic costs and thus unlock alternative routes of adaptation. We additionally hypothesised that a strain which is able to migrate prior to mutation would also ease starvation-induced selection pressures and could facilitate yet more mutational routes. For this experiment we therefore utilised an additional immotile variant of SBW25, which unlike AR2 did not have a transposon inserted into *viscB* (see materials and methods) and thus could migrate via a form of sliding motility prior to mutation (SBW25-Δ*fleQ* (herafter Δ*fleQ*), Alsohim et al. 2014). We observed a ‘blebbing’ phenotype (fig. 1A) in Δ*fleQ* lines despite their ability to migrate in a dendritic fashion; however, we also found blebbing was less frequent under richer nutrient regimes (where populations migrated more rapidly utilising viscosin, see materials and methods). Overall, there was no evidence that the prevalence of the mutational hotspot *ntrB* A289C changed with nutrient condition (Gene-by-environment interaction: χ^2^= 0.9375, df = 7, *P* = 0.9958, see fig. 2). Instead, we observed that the *ntrB* A289C mutation was robust across all tested conditions, featuring in 90-100% of the Δ*fleQ* strains and 80-100% of AR2 strains (fig. 2).

**Fig. 2.**
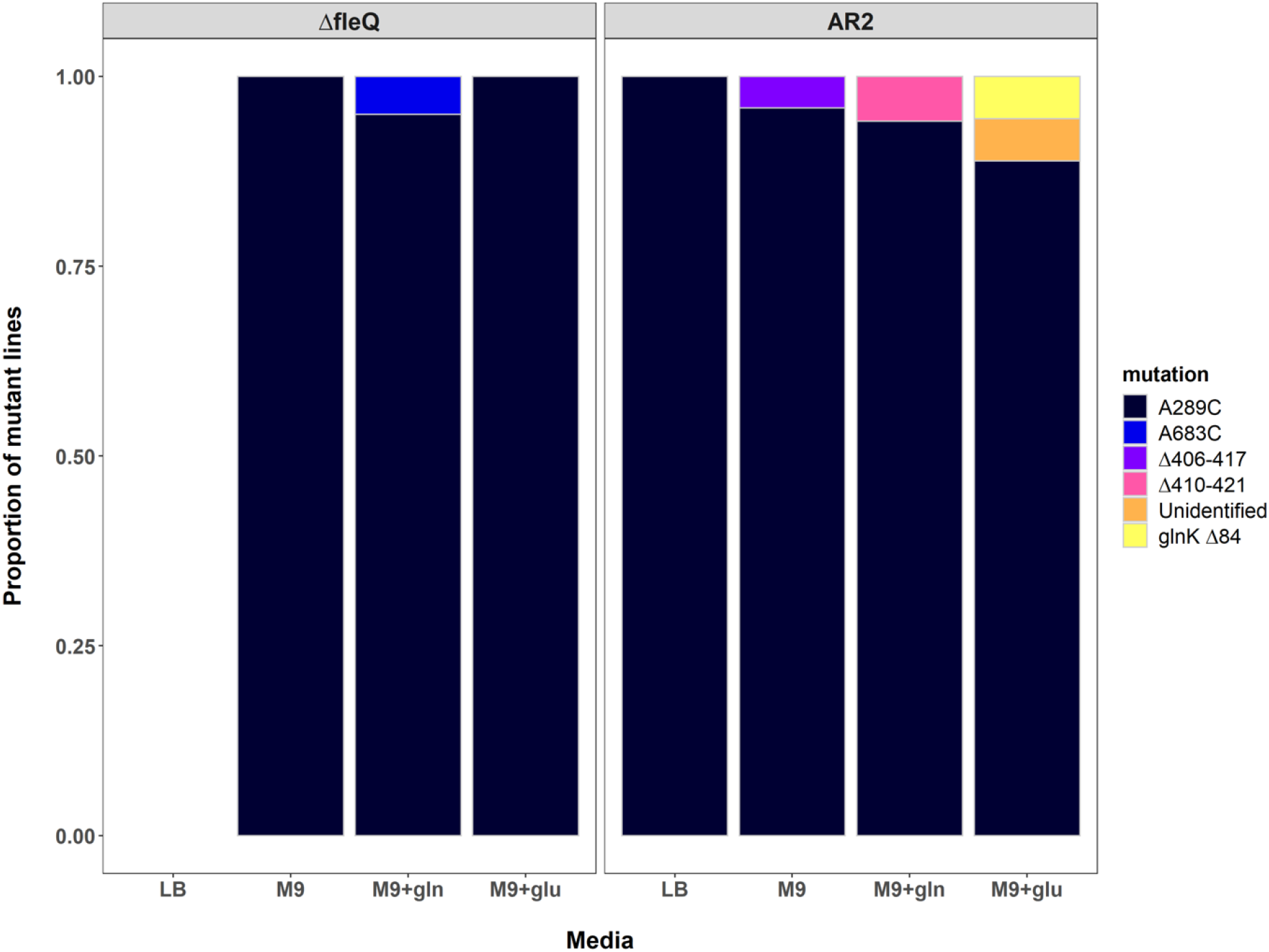
Repeatability of the A289C *ntrB* mutation across genetic background and nutrient environment (total *N* = 116). The proportion of each observed mutation is shown on the y axis. *ntrB* mutation A289C was robust across both strain backgrounds (SBW25Δ*fleQ* shown as Δ*fleQ*, and AR2) and the four tested nutritional environments, remaining the primary target of mutation in all cases (>87%). Lines were evolved using 4mM, 8mM and 16mM of amino acid supplement (see materials and methods). No significant relationship between supplement concentration and evolutionary target was observed (Kruskal-Wallis chi-squared tests: AR2 M9+glu, df = 2, *P* > 0.2; AR2 M9+gln, df = 1, *P* > 0.23; Δ*fleQ* M9+gln, df = 1, *P* > 0.3), as such they are treated as independent treatments for statistical analysis but visually grouped here for convenience. Δ*fleQ* lines evolved on LB were able to migrate rapidly through sliding motility alone, masking any potential emergent flagellate blebs (see Alsohim et al., 2014). Sample sizes (*N*) for other categorical variables: Δ*fleQ* – M9: 25, M9+gln: 20, M9+glu: 7; AR2 −LB: 5, M9: 24, M9+gln: 17, M9+glu: 18.

### Repeatable evolution occurs despite motility being accessible via several mutational routes

Our evolution experiments across nutrient regimes uncovered three novel mutational routes that were observed in a small number of mutants (fig. 2), revealing that mutational accessibility could not explain the level of observed parallel evolution. Most notably was a non-synonymous A-C transversion mutation at site 683 (*ntrB* A683C) in a Δ*fleQ* line evolved on M9+gln, resulting in a missense mutation within the NtrB histidine kinase domain. As a single A-C transversion within the same locus, we may expect A683C to mutate at a similar rate to A289C. We also observed a 12 base-pair deletion from sites 410-421 (*ntrB* Δ410-421) in an AR2 line evolved on M9+gln. Furthermore, we discovered a double mutant in an AR2 line evolved on M9+glu: one mutation was a single nucleotide deletion at site 84 within *glnK*, and the second was another A to C transversion at site 688 resulting in a T230P missense mutation within RNA polymerase sigma factor 54.

GlnK is NtrB’s native regulatory binding partner and repressor in the ntr pathway, meaning the frameshift mutation alone likely explains the observed motility phenotype. However, as this mutant underwent two independent mutations we will not consider it for the following analysis. In addition, *ntrB* Δ410-421 and *ntrB* Δ406-417, despite targeting different nucleotides, translate into identical protein products (both compress residues LVRGL at positions 136-140 to a single L at position 136). Therefore, we will also group them for the following analysis. Under the assumptions that the three remaining one-step observed mutational routes to novel proteins are (*i*) equally likely to appear in the population and (*ii*) equally likely to reach fixation, the original observation of *ntrB* A289C appearing in 23/24 cases becomes exceptional (Bootstrap test: n = 1000000, *P* < 1 x 10^−6^). The likelihood of our observing this by chance, therefore, is highly unlikely. This means that one or both assumptions are almost certainly incorrect. Either the motility phenotype facilitated by the mutations may be unequal, leading to fixation bias. Or the mutations may appear in the population at different rates, resulting in mutation bias. One or both of these elements must be skewing evolution to such a degree that parallel evolution to nucleotide resolution becomes highly predictable.

### Fixation bias cannot explain repeatability to nucleotide resolution

The Darwinian explanation for parallel evolution is that the observed mutational path is outcompeting all others on their way to fixation. If selection alone was driving repeatable evolutionary outcomes, the superior fitness of the *ntrB* A289C genotype should have allowed it to out-migrate other motile genotypes co-existing in the population. To test if the *ntrB* A289C mutation granted the fittest motility phenotype, we allowed the evolved genotypes (A289C, Δ406-417, A683C and *glnK* Δ84) to migrate independently on the four nutritional backgrounds and measured their migration area after 48 h. To allow direct comparison, we first engineered the *ntrB* A683C mutation, which originally evolved in the *ΔfleQ* background, into an AR2 strain. We observed that the non-*ntrB* double mutant, *glnK* Δ84, migrated significantly more slowly than *ntrB* A289C in all four nutrient backgrounds (M9: *P* = 0.00153, M9+gln: *P* = 0.0229, M9+glu: *P* = 0.00460, LB: *P* = 0.00476, fig. 3A). However, *ntrB* A289C did not significantly outperform either of the alternative *ntrB* mutant lines in any environmental condition (*P* value range = 0.0567 – 0.878 fig. 3A). This suggests that selection may have played a role in driving parallel evolution to the level of the *ntrB* locus, but it cannot explain why nucleotide site 289 was so frequently radiated.

**Fig. 3.**
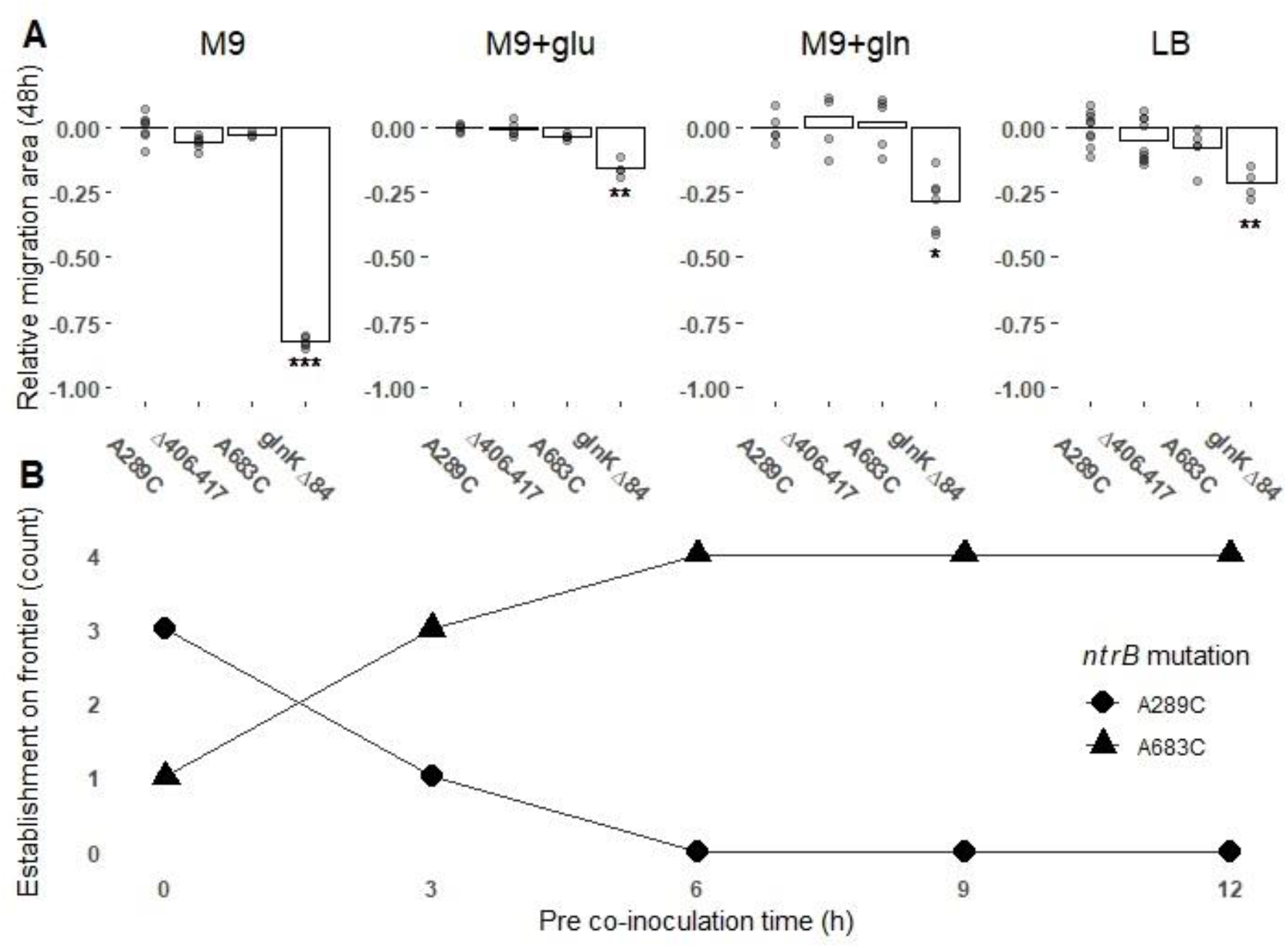
Selection does not strongly favour *ntrB* A289C motility over alternative *ntrB* mutations. (A) Surface area of motile zones following 48h of growth across four environmental conditions. Individual data points from biological replicates are plotted and each migration area has been standardised against the surface area of a *ntrB* A289C mutant grown in the same environment (*ntrB* A289C growth mean = 0). Significance values: * = *P* < 0.05, ** = *P* < 0.01 (Kruskal-Wallis post-hoc Dunn test). (B) *ntrB* A289C lines fail to reach the growth frontier within 6 h of competitor pre-inoculation. Two *ntrB* mutant lines, A289C and A683C, were co-inoculated in equal amounts on soft agar, either immediately (0 h) or with A289C being added at 3 h time points up to 12 h (x-axis) into the centre of an A683C inoculated zone. The strains were competed for 24 h prior to sampling from the motile zone’s leading edge. Genotype establishment at the frontier across the four replicates is shown on the y-axis with the number of lineages at the leading edge represented as 0-4.

To determine if this result remained true when mutant lines were competing in the same population, we directly competed *ntrB* A289C against *ntrB* A683C on M9 minimal medium. In brief, we co-inoculated the two mutant lines on the same soft agar surface and allowed them to competitively migrate before sampling from the leading edge after 24 h of competition. The length of competition was maintained throughout the assay, but *ntrB* A683C lines were allowed to migrate for between 0 and 12 h before the addition of *ntrB* A289C to the agar. We observed that *ntrB* A289C was found predominantly on the leading edge (3/4 replicates) when the mutants were inoculated concurrently, but invading populations of the common genotype swiftly became unable to establish themselves at the leading edge within a narrow time window of 3 h (fig. 3B). This result highlights that in minimal medium *ntrB* A289C does offer a slight dominant phenotype, but to ensure establishment at the leading edge the genotype would need to appear in the population within a handful of generations of a competitor. Given that the range in time before a motility phenotype was observed could vary considerably between independent lines (fig. 1B), our data do not support the hypothesis that global mutation rate could be high enough to allow multiple phenotype-granting mutations to appear in the population almost simultaneously. More likely is that each independent line adhered to the “ early bird gets the worm” maxim, i.e. the *ntrB* mutant which was the first to appear in the population was the genotype that reached fixation. This therefore suggests that the reason *ntrB* A289C is so frequently collected when sampling is due to an evolutionary force other than selection and mutational accessibility.

### Silent genetic variation can break a mutational hotspot

Local mutational biases can play a key role in evolution (Bailey et al. 2017; Lind et al. 2019). Such biases can be introduced by changing DNA curvature (Duan et al. 2018) or through neighbouring tracts of reverse-complement repeats (palindromes and quasi-palindromes), which have been shown to invoke local mutation biases by facilitating the formation of single-stranded DNA hairpins (De Boer and Ripley 1984). Therefore we next searched for a local mutation bias at *ntrB* site 289. Previously, we re-evolved motility in two engineered immotile strains of *P. fluorescens*, AR2 (derived from SBW25) and Pf0-2x (derived from Pf0-1) (Taylor et al. 2015). Although evolved lines in AR2 frequently targeted *ntrB*, Pf0-2x lines fixed mutations across the ntr regulatory pathway. Furthermore, although Pf0-2x did acquire *ntrB* mutations in multiple independent lines, we observed no evidence of *ntrB* site 289 being targeted (Taylor et al. 2015). The NtrB proteins of SBW25 and Pf0-1 are highly homologous (95.57% identity) but share less identity at the genetic level (88.88% identity). A considerable portion of this genetic variation is explained by synonymous genetic variation (8.34%) rather than non-synonymous variance (2.76%). Synonymous mutations can play a role in altering local mutation rates. This may occur by altering the nucleotide-triplet to one with a higher mutation rate (Long et al. 2014) or by altering the secondary structure of longer DNA tracts via the mechanisms outlined above. Nucleotides that remain unpaired when their neighbouring nucleotides form hairpins with nearby reverse-complement tracts have been observed to exhibit increased mutation rates (Wright et al. 2003). Both SBW25 and Pf0-1 were found to have short reverse-complement tracts that flanked site 289, however the called hairpins were not entirely identical in their composition owing to synonymous variance (supplementary fig. S2). Overall, there are 6 synonymous nucleotide substitutions ± 5 codons flanking site 289 (C276G, C279T, C285G, C291G, T294G and G300C), which may have been affecting such hairpin formations and impacting local mutation rate.

To test if synonymous sequence was biasing evolutionary outcomes, we replaced the 6 synonymous sites in an AR2 strain with those from a Pf0-1 background (hereafter AR2-sm). Not all these sites formed part of a theoretically predicted stem that overlapped with site 289, but all were targeted due to their close proximity with the site. AR2-sm lines were placed under selection for motility and we observed that these lines evolved motility significantly more slowly (fig. 4A), both in M9 minimal medium and LB (Wilcoxon rank sum tests with continuity correction: M9, W = 44.5, *P* < 0.001; LB, W = 22, *P* < 0.001). Evolved AR2-sm lines that re-evolved motility within 8 days were sampled and their *ntrB* locus analysed by Sanger sequencing (fig. 4B). We observed some similar *ntrB* mutations to those identified previously: the *ntrB* A683C mutation was observed in one independent line evolved on LB, and *ntrB* Δ406-417 was also observed in both strain backgrounds. However, the most common genotype of *ntrB* A289C fell from being observed in over 95% of independent lines in M9 to 0%. Furthermore, we observed multiple previously unseen *ntrB* mutations, while a considerable number of lines reported wildtype *ntrB* sequences, instead either targeting another gene of the ntr pathway (*glnK*) or unidentified targets that may lay outside of the network (fig. 4B).

**Fig. 4.**
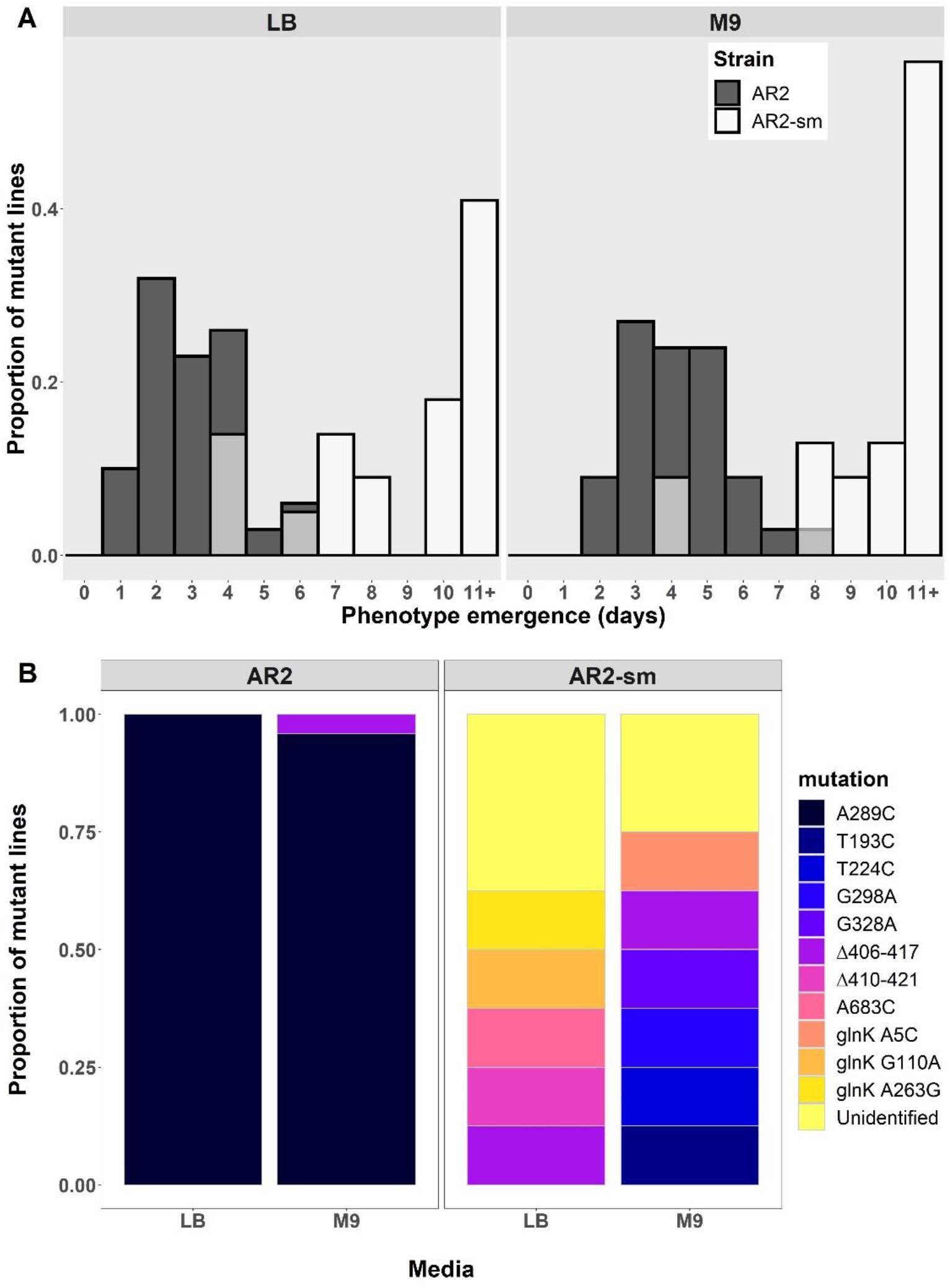
Loss of repeatable evolution conferred by a synonymous sequence mutant (AR2-sm). (A) Histogram of motility phenotype emergence times across independent replicates of immotile SBW25 (AR2) and an AR2 strain with 6 synonymous substitutions in the *ntrB* locus (AR2-sm) in two nutrient conditions. (B) Observed mutational targets across two environments (AR2: LB *N* = 5, M9 *N* = 24; AR2-sm: LB *N* = 8, M9 *N* = 8). Note that characterised genotypes were sampled within 8 days of experiment start date. Unidentified mutations could not be distinguished from wild type sequences of genes belonging to the nitrogen regulatory pathway *(ntrB, glnK* and *glnA*) which were analysed by Sanger sequencing (supplementary table. S1). *ntrB* Δ406-417 was the only mutational target shared by both lines within the same nutritional environment.

To test that the A289C transversion remained a viable mutational target in the AR2-sm genetic background, we subsequently engineered the AR2-sm strain with this motility-enabling mutation. We observed that AR2-sm *ntrB* A289C was motile and comparable in phenotype to a *ntrB* A289C mutant that had evolved in the ancestral AR2 genetic background (supplementary fig. S3). We additionally found that AR2-sm *ntrB* A289C retained comparable motility to the other *ntrB* mutants evolved from AR2-sm (supplementary fig. S3). Therefore, we can determine that the AR2-sm genetic background would not prevent motility following mutation at *ntrB* site 289, nor does it render such a mutation uncompetitive. This therefore infers that the sole variable altered between the two strains (the 6 synonymous changes) are precluding radiation at site 289. Taken together these results strongly suggest that the synonymous sequence immediately surrounding *ntrB* site 289 facilitates its position as a local mutational hotspot, and that local mutational bias is imperative for realising extreme parallel evolution in our model system.

### Silent variation can build a mutational hotspot

As the previous result exemplified the power of synonymous variation in breaking mutational hotspots, we next hypothesised that the same amount of variation could just as readily build a mutational hotspot. To achieve this we engineered a synonymous variant of the immotile Pf0-2x strain (Pf0-2x-sm6). This strain was a reciprocal mutant of AR2-sm, in that it had synonymous variations at the same six sites within *ntrB* but substituted so that they matched AR2’s native sequence (G276C, T279C, G285C, G291C, G294T and C300G). We placed both Pf0-2x and Pf0-2x-sm6 under directional selection for motility and observed that Pf0-2x evolved motility slower than Pf0-2x-sm6 (fig. 5A) and targeted a multitude of sites across multiple loci (fig. 5B). In stark contrast, Pf0-2x-sm6 evolved both more quickly (fig. 5A; Wilcoxon rank sum tests with continuity correction: M9, W = 239.5, *P* < 0.001; LB, W = 461.5, *P* < 0.001) and massively more parallel than its native counterpart. Pf0-2x-sm6 fixed *ntrB* A289C in 80% of instances in M9 (8/10 independent lines), despite this *de novo* mutation not appearing once in a Pf0-2x evolved line (0/22 independent lines, fig. 5B). The striking differences between the two strains from a Pf0-2x genetic background (fig. 5) clearly mirror the results observed in the AR2 genetic background (fig. 4). This reveals that a small number of synonymous variations can heavily bias mutational outcomes across genetic backgrounds and between homologous strains.

**Fig. 5.**
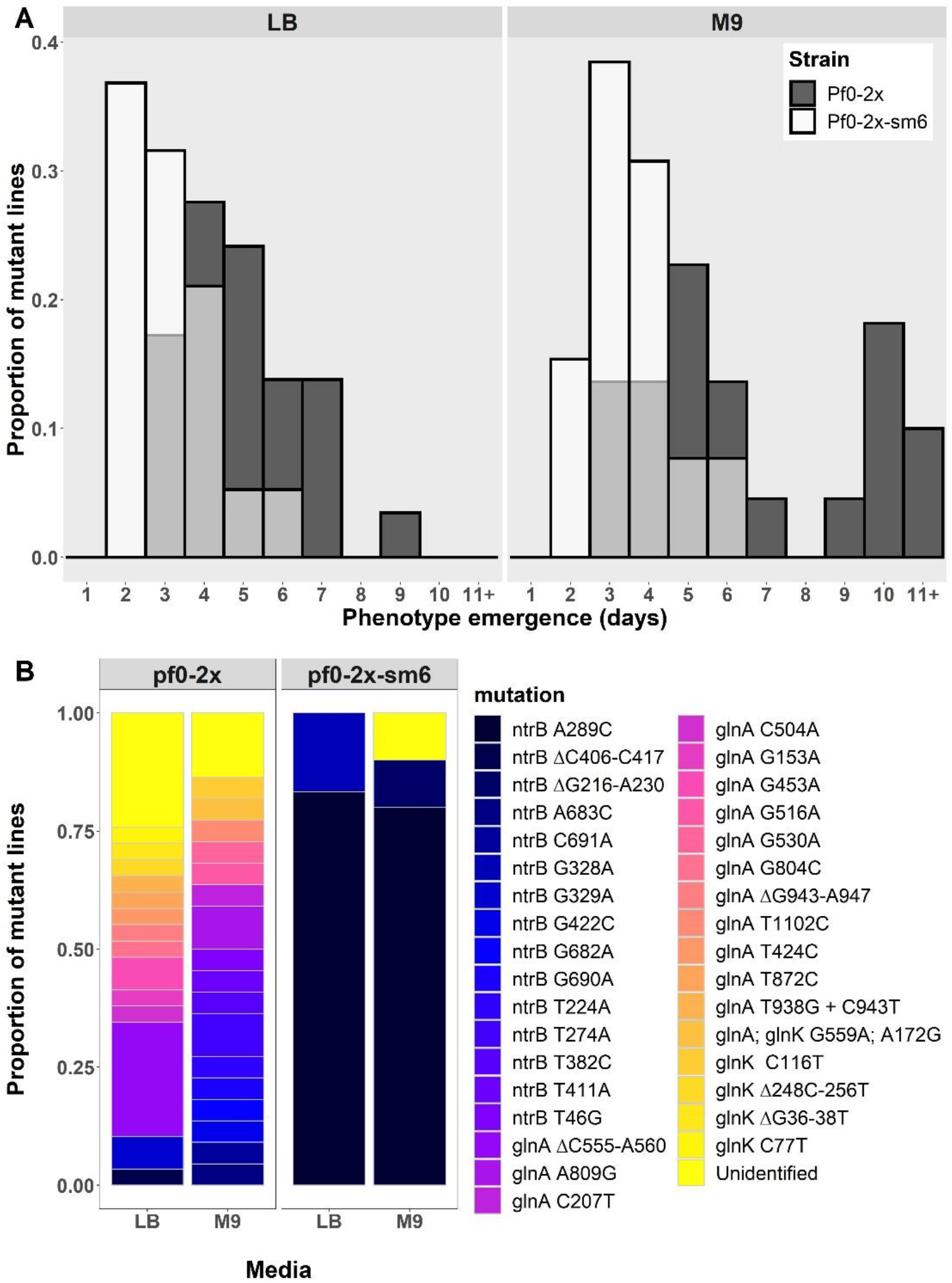
Gain of repeatable evolution conferred by a synonymous sequence mutant (Pf0-2x-sm). (A) Histogram of motility phenotype emergence times across independent replicates of an immotile variant of *P. fluorescens* strain Pf0-1 (Pf0-2x; Taylor et al. 2015) and a Pf0-2x strain with 6 synonymous substitutions in the *ntrB* locus (Pf0-2x-sm) in two nutrient conditions. (B) Observed mutational targets across two environments (Pf0-2x: LB *N* = 29, M9 *N* = 22; Pf0-2x-sm: LB *N* = 6, M9 *N* = 10). Unidentified mutations could not be distinguished from wild type sequences of genes belonging to the nitrogen regulatory pathway *(ntrB, glnK* and *glnA*) which were analysed by Sanger sequencing (supplementary table. S1). Mutation *ntrB A289C* was not observed in a single instance in evolved Pf0-2x lines but became the strongly preferred target following synonymous substitution.

## Discussion

Understanding the evolutionary forces that forge mutational hotspots and repeatedly drive certain mutations to fixation remains an immense challenge. This is true even in simple systems such as the one employed in this study, where clonal bacterial populations were evolved under strong directional selection for very few phenotypes, namely motility and nitrogen metabolism. Here we took immotile variants of *P. fluorescens* SBW25 (AR2) and Pf0-1 (Pf0-2x) that had been observed to repeatedly target the same gene regulatory pathway during the re-evolution of motility (Taylor et al. 2015). We found that evolving populations of AR2 adapted via *de novo* substitution mutation in the same locus (*ntrB*) and at the same nucleotide site (A289C) in over 95% of cases in M9 minimal medium. AR2 populations were constrained in which genetic avenues they could take to access the phenotype under selection, but mutational accessibility and fitness differences alone could not explain such a high degree of parallel evolution. Pf0-2x was distinct in that it did not evolve in parallel to nucleotide nor locus resolution. We observed that by introducing synonymous changes around the mutational hotspot (*ntrB* site 289) in both AR2 and Pf0-2x so that their local genetic sequences were swapped, we could push evolving AR2 populations away from the parallel path and pull Pf0-2x lines onto the parallel path. This work reveals that synonymous sequence is an integral factor toward realising highly repeatable evolution and building a mutational hotspot in our system.

More recent studies have revealed that synonymous changes have an underestimated effect on fitness through their perturbances before and during translation. Synonymous sequence variance can impact fitness by changing the stability of mRNA (Kudla et al. 2009; Kristofich et al. 2018; Lebeuf-Taylor et al. 2019) and altering codons to perturb or better match the codon-anticodon ratio (Frumkin et al. 2018). To our knowledge, we have shown here for the first time that synonymous sequence can also be essential for ensuring parallel evolutionary outcomes across genetic backgrounds. Our results strongly infer that this is due to its impact on local mutational biases, which mechanistically may be owed to the formation of single-stranded hairpins that form between short inverted repeats on the same DNA strand (De Boer and Ripley 1984; Fieldhouse and Golding 1991). The formation of these secondary DNA structures provides a mechanism for intra-locus mutation bias that can operate with extremely local impact and is contingent on DNA sequence variation, as introducing synonymous changes could readily perturb the complementarity of neighbouring inverse repeats (e.g. supplementary fig. S2). Furthermore, the finding of just six synonymous mutations having a significant impact on DNA structure would not represent a surprising result, as secondary structures can be altered by single mutations (Dong et al. 2001).

We can confidently assert that the altered mutational bias is owed to an intralocus effect, owing to the six synonymous sites all residing within 14 bases at either flank of site 289. However the full elucidation of the secondary structure and genetic mechanistic features enabling this powerful mutation bias awaits further study. We know that at least a portion of the 6 substituted nucleotide sites are imperative for parallel genetic outcomes, but we do not yet know if other nucleotide features in the local neighbourhood or more broadly e.g. strand orientation (Merrikh and Merrikh 2018) or distance from the origin of replication (Long et al. 2014) may be combining with local sequence to enforce mutational biases. Interestingly, our data suggest that the mutational hotspot typically mutates so quickly as to mask mutations appearing elsewhere and outside of the nitrogen regulatory pathway, which only appear when the hotspot is perturbed (figs. 4 and 5). This therefore presents the opportunity to additionally quantify the difference in mutation rate owed to secondary structure.

Our findings show that the presence of a mutational hotspot was a stronger deterministic evolutionary force in our system than other variables such as nutrient regime, starvation-induced selection and genetic background. We expected the selective environments to hold some influence over evolutionary outcomes (Bailey et al. 2015) mostly owing to varying levels of antagonistic pleiotropy, which has been found to be a key driver in similar motility studies (Fraebel et al. 2017). Similarly, while parallel evolution can sometimes be impressively robust across genetic backgrounds (Vogwill et al. 2014), some innovations are strongly determined by an organism’s evolutionary history (Blount et al. 2012). Genomic variation also typically combines with environmental differences to drive populations down diverse paths (Spor et al. 2014). However in our experiments, the strains that share the same 6 synonymous sites evolve more similarly than those that share the same broader genetic background (figs. 4B and 5B). These results show that strains can share not only high global homology but also similar genomic architecture – including translated protein structures and gene regulatory network organisation – and yet can have strikingly different mutational outcomes when under selection for the exact same traits owing to synonymous variation. This presents intriguing questions as to whether neutral changes could facilitate the dominance of a genotype during adaptation because of a previously acquired mutational hotspot, and asks whether these mutational hotspots can be selectively enforced.

Models looking to describe drivers of adaptive evolution often place precedence on fitness and the number of accessible adaptive routes (Orr 2005; Krug 2019) yet pay little attention to local mutational biases (however, see Sackman *et al*., 2017). However, mutation rate heterogeneity becomes of paramount importance when systems adhere to the Strong Selection Weak Mutation model (SSWM), which describes instances when an advantageous mutation undergoes a hard sweep to fixation before another beneficial mutation appears (Gillespie 1984). In such cases relative fitness values between adaptive genotypes are relegated to secondary importance behind the likelihood of a genotype appearing in the population. Indeed, experimental systems that adhere to the SSWM maxim have been observed to evolve in parallel despite the option of multiple mutational routes to improved fitness (Vogwill et al. 2014). This suggests that uneven mutational biases can be a key driver in forming mutational hotspots and realising parallel evolution, a conclusion which has been reinforced theoretically (Bailey et al. 2017) although empirical data is still lacking. Understanding the mechanistic causes of mutation rate heterogeneity, therefore, will be essential if we are to determine the presence of mutational hotspots that allow for accurate predictions of evolution (Bailey et al. 2018; Lind et al. 2019). The challenge remains in identifying what these mechanistic quirks may be, where they may be found, and determining how they impact evolutionary outcomes.

Our work sheds light on the ability of silent genetic variation to build a mutational hotspot with functionally significant evolutionary outcomes. This hotspot is built by an adaptive site under strong directional selection that enjoys a biased mutation rate, facilitating highly repeatable evolution when mutation rate and selection align. Mutation is inherently a random process, but not all sites in the genome possess equal fixation potential. Most changes will not improve a phenotype under selection, and those that do will not necessarily mutate at the same rates. Therefore, we can increase our ability to anticipate the location of a mutational hotspot dramatically, permitting we have a detailed understanding of the evolutionary variables at play. Considerable inroads have already been made toward realising this goal. When searching for adaptive targets, it has been highlighted that loss-of-function mutations are the most frequently observed mutational type under selection (Kimura, 1968; Lind, Farr and Rainey, 2015) and that a gene’s wider position within its regulatory network determines its propensity in delivering phenotypic change (McDonald et al. 2009). When searching for mutational biases, it has been shown that parallel evolution at the level of the locus is partially determined by gene length (Bailey et al. 2018) and that molecular apparatus involved in replication and repair can strongly influence the likelihood of a given nucleotide substitution (Lind and Andersson 2008; Stoltzfus and McCandlish 2017). Here, we show that synonymous sequence warrants consideration alongside these other variables by highlighting its impact on the realisation of highly repeatable evolution.

## Materials and Methods

### Model System

Our model system employs strains of the soil microbe *P. fluorescens* SBW25 and Pf0-1 that lack motility through partial gene deletion or disruption of *fleQ*, the master regulator of flagellar motility (Robleto et al. 2003; Alsohim et al. 2014). Motility can be recovered in the absence of *fleQ* following *de novo* mutation that allows for the recruitment of a homologous response regulator, of which the most readily targeted is *ntrC* of the nitrogen regulatory pathway. The initial mutation that facilitates *ntrC* recruitment occurs in other loci in the nitrogen pathway, resulting in the hyper-phosphorylation of *ntrC* (Taylor et al. 2015). Two SBW25-derived strains were used as ancestors in this study: SBW25 Δ*fleQ* (hereafter Δ*fleQ*) and a Δ*fleQ* variant with a functional *viscB* knockout isolated from a transposon library (SBW25Δ*fleQ* IS-ΩKm-hah: PFLU2552, hereafter AR2; Alsohim *et al*., 2014). Δ*fleQ* can migrate on soft agar (0.25%) prior to mutation via a form of sliding motility, which is owed to the strain’s ability to produce viscosin. AR2 cannot produce viscosin and is thus rendered completely immotile prior to mutation. Pf0-1 is a native *gacA* mutant (Seaton et al. 2013) thus does not make viscosin, therefore its Δ*fleQ* variant, Pf0-2x, is rendered completely immotile following disruption of *fleQ*. All cells were grown at 27°C and all strains used throughout the study (ancestral, evolved and engineered) were stored at −80°C in 20% glycerol. The nutrient conditions used throughout the work were lysogeny broth (LB) and M9 minimal media containing glucose and 7.5 mM NH4. The minimal media was used in isolation or supplemented with either glutamate (M9+glu) or glutamine (M9+gln) at a final supplement concentration of 8 mM unless stated otherwise.

### Motility Selection Experiment

Immotile variants were placed under selection for flagella-mediated motility using LB and M9 soft agar (0.25%) motility plates. Details of agar preparation are described in Alsohim et al., 2014. Supplemented concentrations of glutamate (glu)/glutamine (gln) in M9 soft agar were expanded to include final concentrations at 4 mM, 8 mM and 16 mM, as it was observed that biosurfactant-mediated dendritic motility in Δ*fleQ* lines was enhanced at higher supplement concentrations, which masked any emergent blebs (data not shown). Lowering the gln supplement concentration improved the likelihood of observing an emergent flagella bleb in M9+gln motility plates (16 mM: 4/12, 8 mM: 9/20, 4 mM: 7/12 independent lines). However, dendritic motility remained high on all supplements of M9+glu and persistently masked blebbing (16 mM: 2/12, 8 mM: 3/20, 4 mM: 2/11 independent lines). Although gln/glu supplementation had no bearing on motility in AR2 lines, supplement conditions across both gln/glu were expanded for consistency. Single clonal colonies were inoculated into the centre of the agar using a sterile pipette tip and monitored daily until emergence of motile bleb zones (as visualised in fig. 1A). Samples were isolated from the leading edge, selecting for the strongest motility phenotype on the plate, within 24 h of emergence and streaked onto LB agar (1.5%) to obtain a clonal sample. As Δ*fleQ* lines were motile via dendritic movement prior to re-evolving flagella motility and could visually mask flagella-mediated motile zones, samples were left for 120 h prior to sampling from the leading edge of the growth. An exception was made in instances where blebbing motile zones were observed solely further within the growth area, in which case this area was preferentially sampled.

### Sequencing

Motility-facilitating changes were determined through PCR amplification and sequencing of *ntrB, glnK* and *glnA* genes (supplementary table S1). Polymerase chain reaction (PCR) products and plasmids were purified using Monarch® PCR & DNA Cleanup Kit (New England Biolabs) and Sanger sequencing was performed by Eurofins Genomics. A subset of AR2 samples evolved on different nutritional backgrounds was additionally screened through Illumina Whole-Genome Sequencing by the Milner Genomics Centre and MicrobesNG (LB: n = 5, M9: n = 6, M9+gln: n = 6, M9+glu: n = 7). This allowed us to screen for potential secondary mutations and to identify rare changes in motile strains with wildtype *ntrB* sequences. *P. fluorescens* SBW25 genome was used as an assembly template (NCBI Assembly: ASM922v1, GenBank sequence: AM181176.4) and single nucleotide polymorphisms were called using Snippy with default parameters (Seemann 2015) through the Cloud Infrastructure for Microbial Bioinformatics (CLIMB; Connor *et al*., 2016). In instances where coverage at the called site was low (≤10x), called changes were confirmed by Sanger sequencing.

### Soft Agar Motility Assay

Cryopreserved samples of AR2 and derived *ntrB* mutants were streaked and grown for 48 h on LB agar (1.5%). Three colonies were then picked, inoculated in LB broth and grown overnight at an agitation of 180 rpm to create biological triplicates for each sample. Overnight cultures were pelleted via centrifugation, their supernatant withdrawn and the cell pellets re-suspended in phosphate buffer saline (PBS) to a final concentration of OD1 cells/ml. 1 μl of each replicate was inoculated into soft-agar by piercing the top of the agar with the pipette tip and ejecting the culture into the cavity as the tip was withdrawn. Plates were incubated for 48 h and photographed. Diameters of concentric circle growths were calculated laterally and longitudinally, allowing us to calculate an averaged total surface area using A= πr^2^. This process was repeated as several independent lines underwent a second-step mutation (Taylor et al. 2015) within the 48 h assay. This phenotype was readily observable as a blebbing that appeared at the leading edge along a segment of the circumference, distorting the expected concentric circle of a clonal migrating population. As such these plates were discarded from the study. By completing additional sets of biological triplicates, we ensured that each sample had at least three biological replicates for analysis.

### Invasion Assay

OD-corrected biological quadruplets of both *ntrB* mutant lines were prepared as outlined above. For each pair of biological replicates, 1 μl of *ntrB* A683C was first inoculated as outlined above and incubated, followed by *ntrB* A289C’s inoculation into the same cavity after the allotted time had elapsed (0 h, 3 h, 6 h, 9 h and 12 h). When inoculated at 0 h, *ntrB* A289C was added to the plate immediately after *ntrB* A683C. In instances where *ntrB* A289C was added to the plate ≤6 h after *ntrB* A683C, overgrowth of culture was avoided by incubating *ntrB* A289C cultures at 22°C at 0 h until cell pelleting and re-suspension approximately 1 h prior to inoculation. When *ntrB* A289C cultures were added to the plate ≥9 h after *ntrB*-A683C culture, overgrowth of culture was avoided by diluting the culture of *ntrB*-A289C 100-fold into fresh LB broth at 0 h. The same ‘angle of attack’ was used for both instances of inoculation (i.e. the side of the plate that the pipette tip travelled over on its way to the centre), as small volumes of fluid falling from the tip onto the plate could cause local satellite growth. To avoid the risk of satellite growths affecting results, isolated samples were collected from the leading edge 180° from the angle of attack after a period of 24 h. The *ntrB* locus of one sample per replicate was determined by Sanger sequencing to establish the dominant genotype at the growth frontier.

### Genetic engineering

A pTS1 plasmid containing *ntrB* A683C was assembled using overlap extension PCR (oePCR) cloning (for detailed protocol see Bryksin and Matsumura, 2010) using vector pTS1 as a template. The *ntrB* synonymous mutants (AR2-sm and Pf0-2x-sm6) and AR2-sm *ntrB* A289C pTS1 plasmids were constructed using oePCR to assemble the insert sequence for allelic exchange, followed by amplification using nested primers and annealed into a pTS1 vector through restriction-ligation (for full primer list see supplementary table. S1). pTS1 is a suicide vector, able to replicate in *E. coli* but not *Pseudomonas*, and contains a tetracycline resistance cassette as well as an open reading frame encoding SacB. Cloned plasmids were introduced to *P. fluorescens* SBW25 strains via puddle mating conjugation with an auxotrophic *E. coli* donor strain ST18. Mutations were incorporated into the genome through two-step allelic exchange, using a method outline by Hmelo *et al*., 2015, with the following adjustments: (*i*) *P. fluorescens* cells were grown at 27°C. (*ii*) An additional passage step was introduced prior to merodiploid selection, whereby colonies consisting of *P. fluorescens* cells that had incorporated the plasmid (merodiploids) were allowed to grow overnight in LB broth free from selection, granting extra generational time for expulsion of the plasmid from the genome. (*iii*) The overnight cultures were subsequently serially diluted and spot plated onto NSLB agar + 15% (wt/vol) sucrose for AR2 strains and NSLB agar + 5% (wt/vol) sucrose for the Pf0-2x strain. Positive mutant strains were identified through targeted Sanger sequencing of the *ntrB* locus. Merodiploids, which have gone through just one recombination event, will possess both mutant and wild type alleles of the target locus, as well as the *sacB* locus and a tetracycline resistance cassette. However the wild type allele, *sacB* and tetracycline resistance will be subsequently lost following successful two-step recombination. We therefore also screened these mutant strains for counter-selection escape through PCR-amplification and sequencing of the *sacB* locus and growth on tetracycline. Mutants were only considered successful if there was no product on an agarose gel following amplification of *sacB* alongside appropriate controls, the lines were sensitive to tetracycline, and PCR results of the target locus reported expected changes at the targeted sites.

### Statistics

All statistical tests and figures were produced in R (R Core Team 2014). Figures were created using the *ggplot* package (Wickham 2016). A simulated dataset was produced for the Bootstrap test by randomly drawing from a pool of 3 values with equal weights 24 times for 1 million iterations. Note that as the simulated dataset draws from a pool of 3 values, it encodes that no other mutational routes are possible aside from the observed 3. As such the derived statistic is an underestimate, with additional routes at any weight lowering the likelihood of repeat observations of a single value. All other tests were completed using functions in base-R aside from the Dunn test, which was performed using the *FSA* package (Ogle et al. 2020). Along with the Bootstrap test, the statistical tests used throughout the study were: Kruskal-Wallis chi-squared tests, Kruskal-Wallis post-hoc Dunn test, and Wilcoxon rank sum tests with continuity correction.

## Data availability

All raw data used for generation of this manuscript is publicly available and can be accessed at https://github.com/J-S-Horton/Syn-sequence-parallel-evolution.

## Acknowledgments and funding information

We thank Laurence Hurst for comments on earlier versions of this manuscript. In addition we thank member of the Taylor lab Matthew Shepherd for insightful comments and discussion, and Mark Silby for contributing the ancestral *P. fluorescens* Pf0-2x strain used in the study. This work was supported by the University of Bath University Research Studentship Account (URSA) awarded to TBT and NKP; a Royal Society Dorothy Hodgkin Research Fellowship awarded to TBT (DH150169); and the JABBS Foundation for RWJ. Bioinformatics analysis of the paper was carried out using the Medical Research Council’s (MRC) Cloud Infrastructure for Microbial Bioinformatics (CLIMB), and Illumina Whole-Genome Sequencing by the Milner Genomics Centre, Bath, UK and MicrobesNG, Birmingham, UK.

## Supplementary materials

### Assessing Pleiotropy via Growth Rate

Cryopreserved samples of AR2 and derived *ntrB* mutants were streaked and grown for 48 h on LB agar (1.5%). Three colonies were then picked, inoculated in LB broth and grown overnight at an agitation of 180 rpm to create biological triplicates for each sample. This process was repeated with an independent batch of biological triplicates on a separate day to produce a total of 6 biological replicates for each sample. Overnight cultures were pelleted via centrifugation, their supernatant withdrawn and the cell pellets re-suspended in phosphate buffer saline (PBS) to a final concentration of OD1 cells/ml. The resuspension was subsequently diluted 100-fold into a 96-well plate (Costar**®**) containing nutrient broth. The plates were analysed in a Multiskan **™** FC Microplate Photometer (Thermo Fisher Scientific) for 24h, with autonomous OD readings every 10 min without agitation. Growth values were determined by calculating area under the curve using the trapezoidal rule (approach outlined in Huang and Pang, 2012). This allowed us to incorporate elements of the pleiotropic consequences to metabolism as well as the benefits of the motile swimming phenotype, including prolonged lag phases, steeper exponential phases and differing eventual yields achieved by mutant populations relative to the ancestral strain (growth curves not shown).

### *ntrB* loci analysis

Theoretical hairpin stem-loop structures were generated using the *mfg* tool and methodology developed by Wright *et al*., 2003. The *mfg* tool is used in conjunction with the Quikfold tool on the DINAMelt Web Server (Markham and Zuker 2005). Default parameters were used for Quikfold with the exception of temperature, which was amended to 27°C. The first 400 nucleotides of the open reading frames of *P. fluorescens* SBW25 *ntrB* and Pf0-1 *ntrB* were used as input sequences, and AR2-sm’s and Pf0-2x-sm’s input sequences were created by manually editing SBW25’s and Pf0-1’s *ntrB* sequence. The *mfg* application generates the most stable stem-loop structure for each base in which the selected base remains unpaired and so is at a higher likelihood of mutation. The window size of neighbouring nucleotides that are used to form the stem-loop structure can be adjusted, and a window length of 40 nucleotides was used for the analysis in this study.

**Fig. S1.**
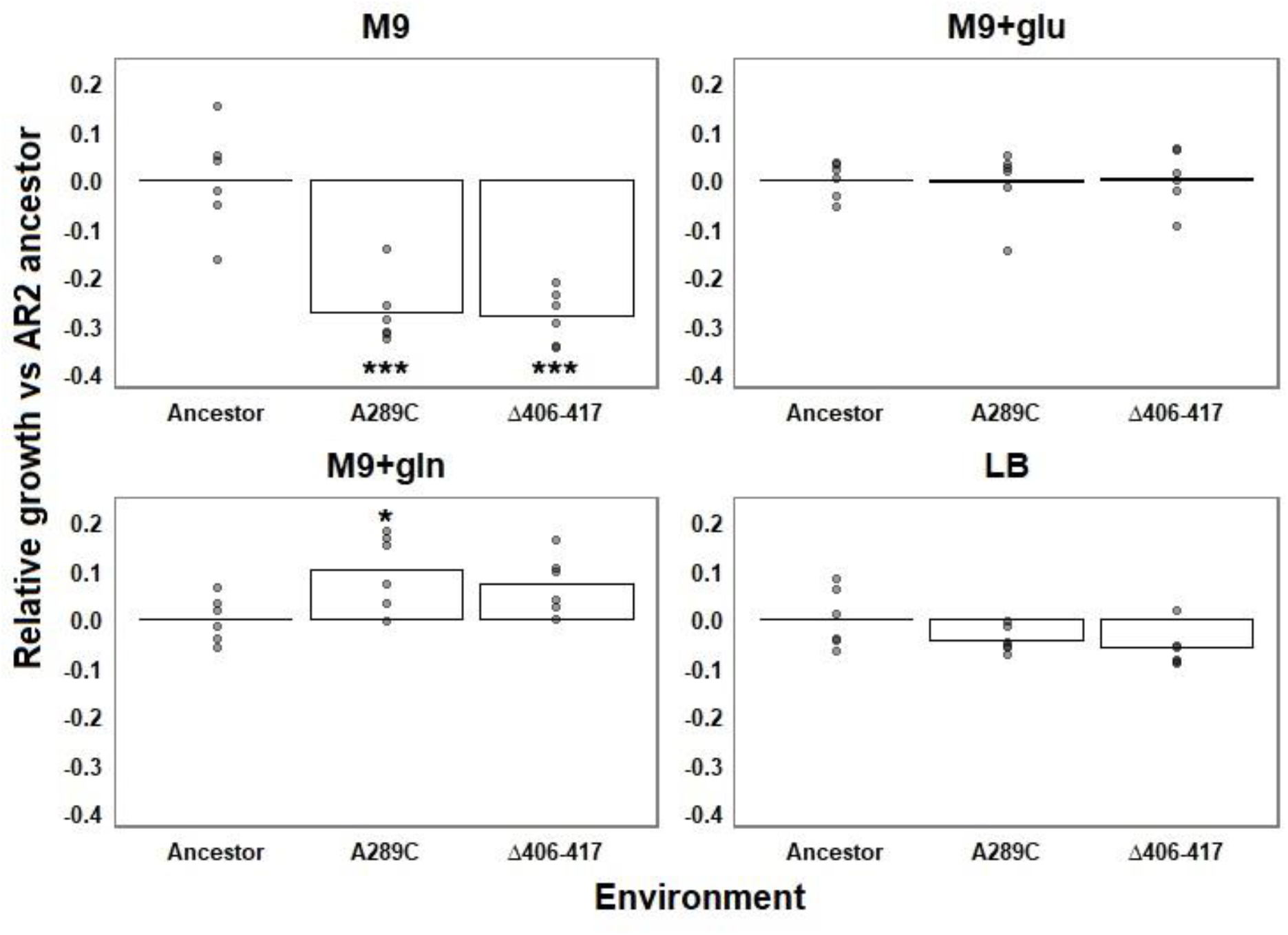
Growth kinetics of mutant AR2 lines in static liquid culture over 24h. Nutrient environments: M9 = M9 minimal media supplemented with NH4 at 7.5 mM. M9+glu = additional glutamate added at 8 mM. M9+gln = additional glutamine added at 8 mM. LB = lysogeny broth. Growth yield was determined using area under the curve, and each yield has been standardised against the yield of the AR2 ancestral strain grown in the same environment (AR2 ancestor growth mean = 0). Individual data points from biological replicates are plotted, and ranges around the mean growth of the ancestral strain are shown in column one of each frame. Plots are the means of six biological replicates. Significance values: * = *P* < 0.05, ** = *P* < 0.01, *** = *P* < 0.001 (one-way ANOVA post-hoc Tukey HSD test).

**Fig. S2.**
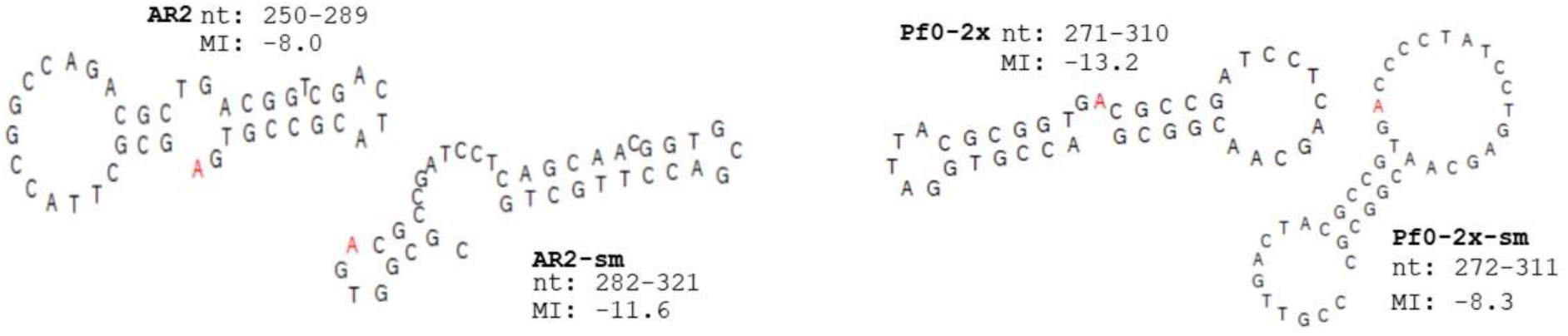
Quasi-palindromic sequences flank *ntrB* site 289 in both *P. fluorescens* SBW25 and Pf0-1 derived strains. Theoretical hairpin formations were generated using the *mfg* program (Wright et al. 2003). This software calculates the most stable hairpin formed between neighbouring tracts (± 40 nucleotides from site 289) in which the site of interest (in this case site 289, highlighted in red) remains unpaired. In these examples the nucleotides are forced into stem-loop structures that have been documented to comprise hairpins (Ripley, 1982). The stability, structure and included nucleotide tracts of the predicted hairpins differ between strains and determine the radiated nucleotide site’s Mutational Index (MI): AR2 = −8.0. AR2-sm = −11.6, Pf0-2x = −13.2, Pf0-2x-sm = −8.3. These differences are partially owed to synonymous sequence variation as highlighted by the altered hairpin formation exhibited by AR2-sm and Pf0-2x-sm, who differ from their ancestors by 6 synonymous substitutions. AR2 and Pf0-2x-sm, the two strains that evolve in a highly parallel manner, share similar features that are absent in the other two strains. Namely their MI’s are similar (−8.0 and −8.3) and the frequently radiated ‘A’ is located two nucleotides away from the base of a singular long, stable stem. As the *mfg* program only calls the most stable hairpin configuration it may miss alternative structures that temporarily form and raise mutation rate, however the tool exemplifies the power of synonymous variance in altering hairpin stability.

**Fig. S3.**
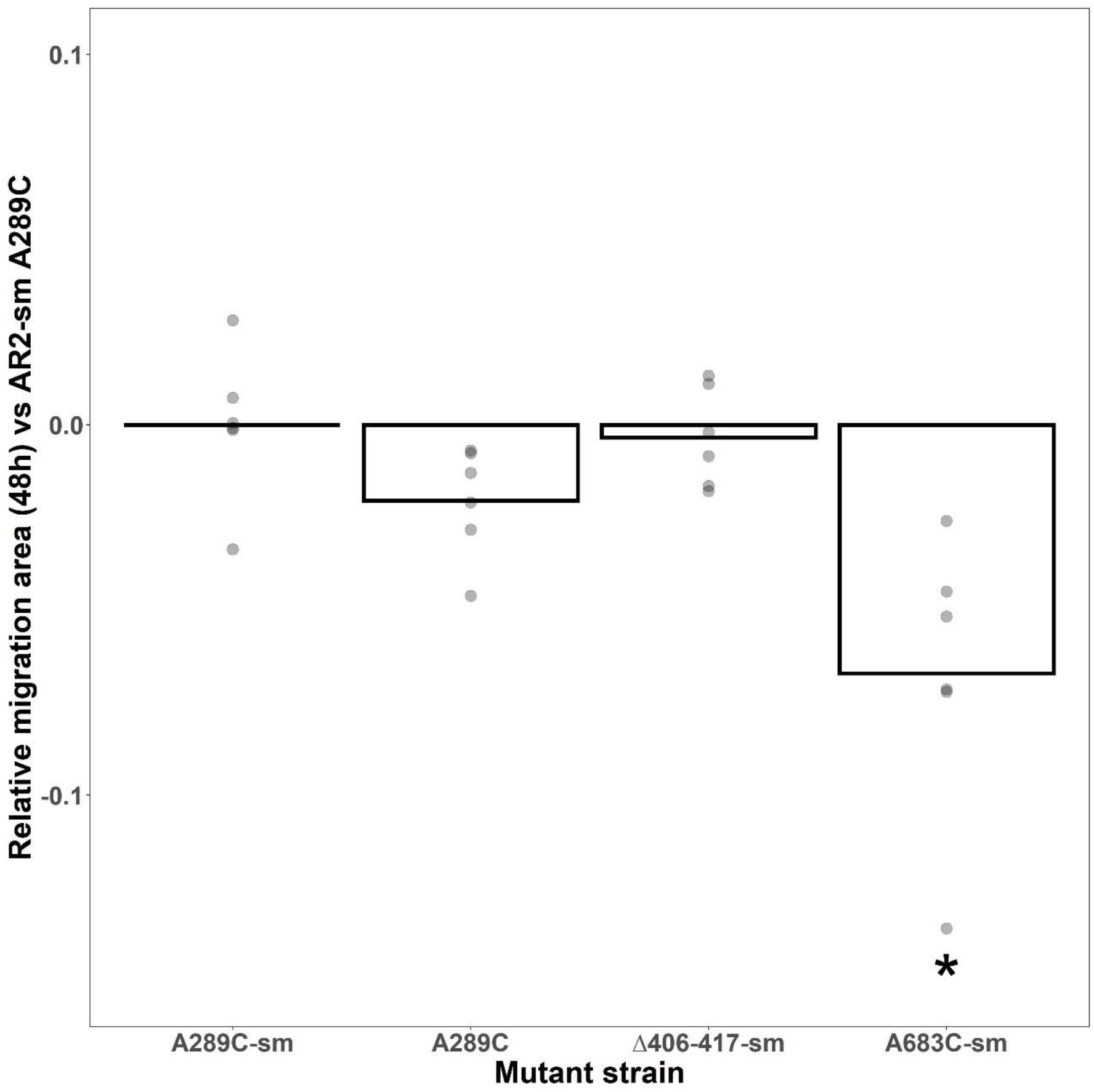
*ntrB* A289C in AR2-sm retains comparative fitness to its ancestral counterpart. The motility phenotype of AR2 *ntrB* A289C and alternative AR2-sm *ntrB* mutants (Δ406-417-sm and A683-sm) were measured against an engineered AR2-sm *ntrB* A289C mutant (A289C-sm) in minimal medium. A289C-sm was not significantly outperformed by any strain, instead showing a significantly superior motility phenotype to A683-sm in M9. Although the two motile lines displayed comparable motility in an AR2 background (fig. 3A), the inferior phenotype observed here may be owed to an uncharacterised secondary mutation. Individual data points from biological replicates are plotted and each migration area has been standardised against the surface area of a *ntrB* A289C-sm mutant grown in the same environment (*ntrB* A289C-sm growth mean = 0). Significance values: * = *P* < 0.05, ** (Kruskal-Wallis post-hoc Dunn test).

**Table. S1.**
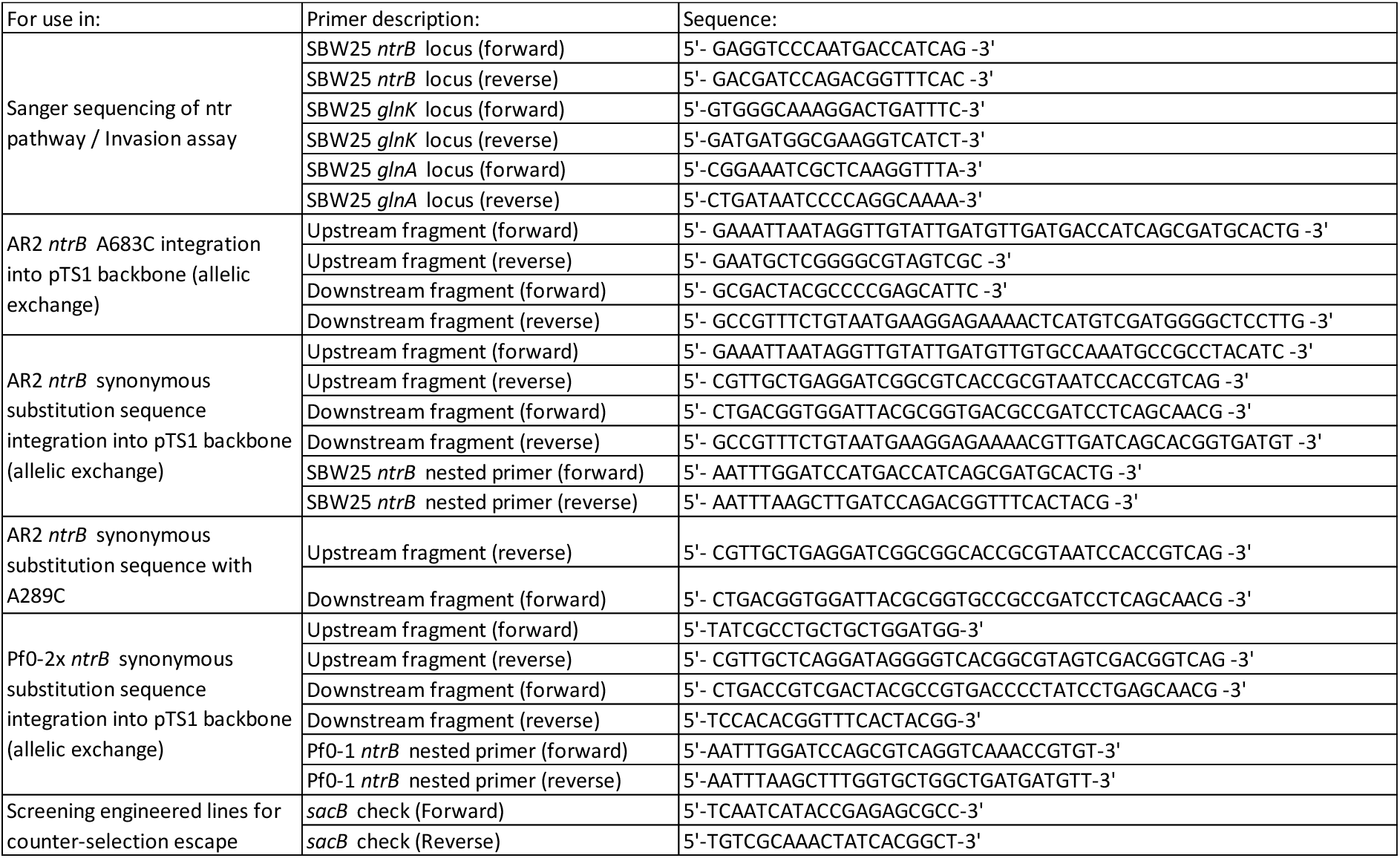
List of primers used throughout the study.

## References

Alsohim AS, Taylor TB, Barrett GA, Gallie J, Zhang X, Altamirano-Junqueira AE, Johnson LJ, Rainey PB, Jackson RW. 2014. The biosurfactant viscosin produced by Pseudomonas fluorescens SBW25 aids spreading motility and plant growth promotion. Environ. Microbiol. 16:2267–2281.

Avrani S, Bolotin E, Katz S, Hershberg R. 2017. Rapid Genetic Adaptation during the First Four Months of Survival under Resource Exhaustion. Mol. Biol. Evol. 34:1758–1769.

Bailey SF, Blanquart F, Bataillon T, Kassen R. 2017. What drives parallel evolution?: How population size and mutational variation contribute to repeated evolution. BioEssays 39:1–9.

Bailey SF, Guo Q, Bataillon T. 2018. Identifying drivers of parallel evolution: A regression model approach. Genome Biol. Evol. 10:2801–2812.

Bailey SF, Rodrigue N, Kassen R. 2015. The effect of selection environment on the probability of parallel evolution. Mol. Biol. Evol. 32:1436–1448.

Barrett RDH, M’Gonigle LK, Otto SP. 2006. The distribution of beneficial mutant effects under strong selection. Genetics 174:2071–2079.

Blount ZD, Barrick JE, Davidson CJ, Lenski RE. 2012. Genomic analysis of a key innovation in an experimental Escherichia coli population. Nature 489:513–518.

De Boer JG, Ripley LS. 1984. Demonstration of the production of frameshift and base-substitution mutations by quasipalindromic DNA sequences.

Bryksin A V, Matsumura I. 2010. Overlap extension PCR cloning: a simple and reliable way to create recombinant plasmids. Biotechniques 48:463–465.

Bull JJ, Badgett MR, Wichman HA, Huehenbeck JP, Hillis DM, Gulati A, Ho C, Molineux IJ. 1997. Exceptional Convergent Evolution in a Virus. Genetics 147:1497–1507.

Connor TR, Loman NJ, Thompson S, Smith A, Southgate J, Poplawski R, Bull MJ, Richardson E, Ismail M, Elwood-Thompson S, et al. 2016. CLIMB (the Cloud Infrastructure for Microbial Bioinformatics): an online resource for the medical microbiology community. Microb. Genomics 2:6.

Van Ditmarsch D, Boyle KE, Sakhtah H, Oyler JE, Carey D, Déziel É, Dietrich LEP, Xavier JB. 2013. Convergent Evolution of Hyperswarming Leads to Impaired Biofilm Formation in Pathogenic Bacteria. Cell Rep 4:697–708.

Dong F, Allawi HT, Anderson T, Neri BP, Lyamichev VI. 2001. Secondary structure prediction and structure-specific sequence analysis of single-stranded DNA. Nucleic Acids Res. 29:3248–3257.

Duan C, Huan Q, Chen X, Wu S, Carey LB, He X, Qian W. 2018. Reduced intrinsic DNA curvature leads to increased mutation rate. Genome Biol. 19:1–12.

Eyre-Walker A, Hurst LD. 2001. The evolution of isochores. Nat. Rev. Genet. 2:549–555.

Fieldhouse D, Golding B. 1991. A source of small repeats in genomic DNA. Genetics 129:563–572.

Fong SS, Joyce AR, Palsson BØ. 2005. Parallel adaptive evolution cultures of Escherichia coli lead to convergent growth phenotypes with different gene expression states. Genome Res.:1365–1372.

Fraebel DT, Mickalide H, Schnitkey D, Merritt J, Kuhlman TE, Kuehn S. 2017. Environment determines evolutionary trajectory in a constrained phenotypic space. Elife 6:e24669.

Frumkin I, Lajoie MJ, Gregg CJ, Hornung G, Church GM, Pilpel Y. 2018. Codon usage of highly expressed genes affects proteome-wide translation efficiency. Proc. Natl. Acad. Sci. U. S. A. 115:E4940–E4949.

Galen SC, Natarajan C, Moriyama H, Weber RE, Fago A, Benham PM, Chavez AN, Cheviron ZA, Storz JF, Witt CC. 2015. Contribution of a mutational hot spot to hemoglobin adaptation in high-Altitude Andean house wrens. Proc. Natl. Acad. Sci. U. S. A. 112:13958–13963.

Gillespie JH. 1984. Molecular Evolution Over the Mutational Landscape. Evolution (N. Y). 38:1116.

Hermisson J, Pennings PS. 2005. Soft sweeps: Molecular population genetics of adaptation from standing genetic variation. Genetics 169:2335–2352.

Herron MD, Doebeli M. 2013. Parallel Evolutionary Dynamics of Adaptive Diversification in Escherichia coli. PLoS Biol. 11:e1001490.

Hmelo LR, Borlee BR, Almblad H, Love ME, Randall TE, Tseng BS, Lin CY, Irie Y, Storek KM, Yang JJ, et al. 2015. Precision-engineering the Pseudomonas aeruginosa genome with two-step allelic exchange. Nat. Protoc. 10:1820–1841.

Huang S, Pang L. 2012. Comparing statistical methods for quantifying drug sensitivity based on in vitro dose-response assays. Assay Drug Dev. Technol. 10:88–96.

Kram KE, Geiger C, Ismail WM, Lee H, Tang H, Foster PL, Finkel SE. 2017. Adaptation of Escherichia coli to Long-Term Serial Passage in Complex Medium: Evidence of Parallel Evolution. mSystems 2:1–12.

Kristofich J, Morgenthaler AB, Kinney WR, Ebmeier CC, Snyder DJ, Old WM, Cooper VS, Copley SD. 2018. Synonymous mutations make dramatic contributions to fitness when growth is limited by a weak-link enzyme.Matic I, editor. PLOS Genet. [Internet] 14:e1007615. Available from: https://dx.plos.org/10.1371/journal.pgen.1007615

Krug J. 2019. Accessibility percolation in random fitness landscapes. bioRxiv [Internet]. Available from: http://arxiv.org/abs/1903.11913

Kudla G, Murray AW, Tollervey D, Plotkin JB. 2009. Coding-sequence determinants of gene expression in Escherichia coli. Science (80-.). [Internet] 324:255–258. Available from: https://www.ncbi.nlm.nih.gov/pmc/articles/PMC3624763/pdf/nihms412728.pdf

Lässig M, Mustonen V, Walczak AM. 2017. Predicting evolution. Nat. Ecol. Evol. [Internet] 1:1–9. Available from: http://dx.doi.org/10.1038/s41559-017-0077

Lebeuf-Taylor E, McCloskey N, Bailey SF, Hinz A, Kassen R. 2019. The distribution of fitness effects among synonymous mutations in a gene under selection. Elife [Internet]:e45952. Available from: https://doi.org/10.7554/eLife.45952.001

Lind PA, Andersson DI. 2008. Whole-genome mutational biases in bacteria. Proc. Natl. Acad. Sci. U. S. A. [Internet] 105:17878–17883. Available from: www.pnas.org/cgi/content/full/

Lind PA, Farr AD, Rainey PB. 2015. Experimental evolution reveals hidden diversity in evolutionary pathways. Elife 4.

Lind PA, Libby E, Herzog J, Rainey PB. 2019. Predicting mutational routes to new adaptive phenotypes. Elife [Internet] 8:e38822. Available from: https://www.ncbi.nlm.nih.gov/pubmed/30616716

Long H, Sung W, Miller SF, Ackerman MS, Doak TG, Lynch M. 2014. Mutation rate, spectrum, topology, and context-dependency in the DNA mismatch repair-deficient Pseudomonas fluorescens ATCC948. Genome Biol. Evol. 7:262–271.

M. Kimura. 1968. Evolutionary Rate at the Molecular Level. Nature [Internet] 217:624–626. Available from: https://www-nature-com.remote.library.osaka-u.ac.jp:8443/articles/217624a0.pdf

Markham NR, Zuker M. 2005. DINAMelt web server for nucleic acid melting prediction. Nucleic Acids Res. 33:577–581.

McDonald MJ, Gehrig SM, Meintjes PL, Zhang XX, Rainey PB. 2009. Adaptive divergence in experimental populations of Pseudomonas fluorescens. IV. Genetic constraints guide evolutionary trajectories in a parallel adaptive radiation. Genetics 183:1041–1053.

Mcgee LW, Sackman AM, Morrison AJ, Pierce J, Anisman J, Rokyta DR. 2016. Synergistic Pleiotropy Overrides the Costs of Complexity in Viral Adaptation. Genetics 202:285–295.

McGrath PT, Xu Y, Ailion M, Garrison JL, Butcher RA, Bargmann CI. 2011. Parallel evolution of domesticated Caenorhabditis species targets pheromone receptor genes. Nature 477:321–325.

Merrikh CN, Merrikh H. 2018. Gene inversion potentiates bacterial evolvability and virulence. Nat. Commun. 9:10.

Meyer JR, Dobias DT, Weitz JS, Barrick JE, Quick RT, Lenski RE. 2012. Repeatability and contingency in the evolution of a key innovation in phage lambda. Science (80-.). [Internet] 335:428–432. Available from: http://science.sciencemag.org/

Miller C, Kong J, Tran TT, Arias CA, Saxer G, Shamoo Y. 2013. Adaptation of Enterococcus faecalis to daptomycin reveals an ordered progression to resistance. Antimicrob. Agents Chemother. 57:5373–5383.

Notley-McRobb L, Ferenci T. 1999. Adaptive mgl-regulatory mutations and genetic diversity evolving in glucose-limited Escherichia coli populations. Environ. Microbiol. 1:33–43.

Ogle DH, Wheeler P, Dinno A. 2020. FSA: Fisheries Stock Analysis. Available from: https://github.com/droglenc/FSA

Orr HA. 2005. THE PROBABILITY OF PARALLEL EVOLUTION. Evolution (N. Y). 59:216.

Ostrowski EA, Woods RJ, Lenski RE. 2008. The genetic basis of parallel and divergent phenotypic responses in evolving populations of Escherichia coli. Proc. R. Soc. B Biol. Sci. 275:277–284.

R Core Team. 2014. R: A language and environment for statistical computing. R Found. Stat. Comput. Vienna, Austria [Internet]. Available from: http://www.r-project.org/.

Riehle MM, Bennett AF, Long AD. 2001. Genetic architecture of thermal adaptation in Escherichia coli. Proc. Natl. Acad. Sci. U. S. A. [Internet] 98:525–530. Available from: www.pnas.orgcgidoi10.1073pnas.021448998

Ripley LS. 1982. Model for the participation of quasi-palindromic DNA sequences in frameshift mutation. Proc. Natl. Acad. Sci. U. S. A. 79:4128–4132.

Robleto EA, López-Hernández I, Silby MW, Levy SB. 2003. Genetic analysis of the AdnA regulon in Pseudomonas fluorescens: Nonessential role of flagella in adhesion to sand and biofilm formation. J. Bacteriol. 185:453–460.

Sackman AM, McGee LW, Morrison AJ, Pierce J, Anisman J, Hamilton H, Sanderbeck S, Newman C, Rokyta DR. 2017. Mutation-driven parallel evolution during viral adaptation. Mol. Biol. Evol. 34:3243–3253.

Seaton SC, Silby MW, Levy SB. 2013. Pleiotropic effects of gaca on pseudomonas fluorescens pf0-1 in vitro and in soil. Appl. Environ. Microbiol. 79:5405–5410.

Seemann T. 2015. Snippy: fast bacterial variant calling from NGS reads. Available from: https://github.com/tseemann/snippy

Sekowska A, Wendel S, Fischer EC, Nørholm MHH. 2016. Generation of mutation hotspots in ageing bacterial colonies. Sci. Rep. 6:4–10.

Spor A, Kvitek DJ, Nidelet T, Martin J, Legrand J, Dillmann C, Bourgais A, De Vienne D, Sherlock G, Sicard D. 2014. Phenotypic and genotypic convergences are influenced by historical contingency and environment in yeast. Evolution (N. Y). 68:772–790.

Stoltzfus A, McCandlish DM. 2017. Mutational biases influence parallel adaptation. Mol. Biol. Evol. 34:2163–2172.

Taylor TB, Mulley G, Dills AH, Alsohim AS, McGuffin LJ, Studholme DJ, Silby MW, Brockhurst MA, Johnson LJ, Jackson RW. 2015. Evolutionary resurrection of flagellar motility via rewiring of the nitrogen regulation system. Science (80-.). 347:1014–1017.

Tenaillon O, Rodríguez-Verdugo A, Gaut RL, McDonald P, Bennett AF, Long AD, Gaut BS. 2012. The molecular diversity of adaptive convergence. Science (80-.). 335:457–461.

Trevino V. 2020. HotSpotAnnotations — a database for hotspot mutations and annotations in cancer. Database: 1–8.

Turner CB, Marshall CW, Cooper VS. 2018. Parallel genetic adaptation across environments differing in mode of growth or resource availability. Evol. Lett. 2:355–367.

Vogwill T, Kojadinovic M, Furió V, Maclean RC. 2014. Testing the role of genetic background in parallel evolution using the comparative experimental evolution of antibiotic resistance. Mol. Biol. Evol. 31:3314–3323.

Weber S, Ramirez C, Doerfler W. 2020. Signal hotspot mutations in SARS-CoV-2 genomes evolve as the virus spreads and actively replicates in different parts of the world. Virus Res. [Internet] 289:198170. Available from: https://doi.org/10.1016/j.virusres.2020.198170

Weinreich DM, Delaney NF, De Pristo MA, Hartl DL. 2006. Darwinian Evolution Can Follow Only Very Few Mutational Paths to Fitter Proteins. Science (80-.). 312.

Wichman HA, Badgett MR, Scott LA, Boulianne CM, Bull JJ. 1999. Different trajectories of parallel evolution during viral adaptation. Science (80-.). 285:422–424.

Wickham H. 2016. ggplot2: Elegant Graphics for Data Analysis. :ISBN 978-3-319-24277-4. Available from: https://ggplot2.tidyverse.org

Wood TE, Burke JM, Rieseberg LH. 2005. Parallel genotypic adaptation: When evolution repeats itself. Genetica 123:157–170.

Woods R, Schneider D, Winkworth CL, Riley MA, Lenski RE. 2006. Tests of parallel molecular evolution in a long-term experiment with Escherichia coli. Proc. Natl. Acad. Sci. U. S. A. 103:9107–9112.

Wright BE, Reschke DK, Schmidt KH, Reimers JM, Knight W. 2003. Predicting mutation frequencies in stem-loop structures of derepressed genes: Implications for evolution. Mol. Microbiol. 48:429–441.

